# A novel mutation in *KCNK16* causing a gain-of-function in the TALK-1 potassium channel: a new cause of maturity onset diabetes of the young

**DOI:** 10.1101/2020.02.04.929430

**Authors:** Sarah M Graff, Stephanie R Johnson, Paul J Leo, Prasanna K Dadi, Arya Y Nakhe, Aideen M McInerney-Leo, Mhairi Marshall, Matthew A Brown, David A Jacobson, Emma L Duncan

**Affiliations:** Department of Molecular Physiology and Biophysics, Vanderbilt University, Nashville, Tennessee, USA; Department of Endocrinology, Queensland Children’s Hospital, Stanley Street, South Brisbane, Australia; Translational Genomics Group, Institute of Health and Biomedical Innovation, Faculty of Health, Queensland University of Technology, Translational Research Institute, Princess Alexandra Hospital, Ipswich Rd, Woolloongabba, Australia; Faculty of Medicine, University of Queensland, Herston, Australia; Dermatology Research Centre, Dermatology Research Centre, The University of Queensland, The University of Queensland Diamantina Institute, Brisbane, QLD, Australia; Guy’s and St Thomas’ NHS Foundation Trust and King’s College London NIHR Biomedical Research Centre, King’s College London, United Kingdom; Department of Endocrinology and Diabetes, Royal Brisbane & Women’s Hospital, Butterfield St, Herston, Australia

**Author notes:** These authors contributed equally to this study (Sarah M Graff, Stephanie R Johnson). These authors contributed equally to this study (David A Jacobson, Emma L Duncan). Corresponding Author: Prof. Emma Duncan at the Department of Endocrinology and Diabetes, Royal Brisbane & Women’s Hospital, Butterfield St, Herston, QLD 4029, Australia. Address reprint requests to A/Prof. David Jacobson at the Department of Molecular Physiology and Biophysics, 2213 Garland Avenue, Nashville, TN, 37232,.

**Keywords:** TALK-1, calcium, MODY, diabetes, channelopathy

## Abstract

**Background:** Maturity-onset diabetes of the young (MODY) is a heterogeneous group of monogenic disorders of impaired glucose-stimulated insulin secretion (GSIS). Mechanisms include β-cell K_ATP_ channel dysfunction (e.g., *KCNJ11* (MODY13) or *ABCC8* (MODY12) mutations); however, no other β-cell channelopathies have been identified in MODY.

**Methods:** A four-generation family with autosomal dominant non-obese, non-ketotic antibody-negative diabetes, without mutations in known MODY genes, underwent exome sequencing. Whole-cell and single-channel K^+^ currents, Ca^2+^ handling, and GSIS were determined in cells expressing either mutated or wild-type (WT) protein.

**Results:** We identified a novel non-synonymous genetic mutation in *KCNK16* (NM_001135105: c.341T>C, p.Leu114Pro) segregating with MODY. *KCNK16* is the most abundant and β-cell-restricted K^+^ channel transcript and encodes the two-pore-domain K^+^ channel TALK-1. Whole-cell K^+^ currents in transfected HEK293 cells demonstrated drastic (312-fold increase) gain-of-function with TALK-1 Leu144Pro vs. WT, due to greater single channel activity. Glucose-stimulated cytosolic Ca^2+^ influx was inhibited in mouse islets expressing TALK-1 Leu114Pro (area under the curve [AUC] at 20mM glucose: Leu114Pro 60.1 vs. WT 89.1; *P*=0.030) and less endoplasmic reticulum calcium storage (cyclopiazonic acid-induced release AUC: Leu114Pro 17.5 vs. WT 46.8; *P*=0.008). TALK-1 Leu114Pro significantly blunted GSIS compared to TALK-1 WT in both mouse (52% decrease, *P*=0.039) and human (38% decrease, *P*=0.019) islets.

**Conclusions:** Our data identify a novel MODY-associated gene, *KCNK16*; with a gain-of-function mutation limiting Ca^2+^ influx and GSIS. A gain-of-function common polymorphism in *KCNK16* is associated with type 2 diabetes (T2DM); thus, our findings have therapeutic implications not only for *KCNK16*-associated MODY but also for T2DM.

## INTRODUCTION

Maturity-onset diabetes of the young (MODY) is a rare monogenic cause of familial diabetes. To date, 13 MODY genes have been confirmed, all involved in pancreatic β-cell insulin secretion and all with autosomal dominant transmission^1^. 2-2.5% of pediatric diabetes cases carry pathogenic/likely pathogenic variants in MODY genes^2,3^; however, MODY is often undiagnosed, either because the diagnosis is not considered^4^ or because genetic screening is limited. There are also cases with compelling clinical histories in whom, despite comprehensive screening of known MODY genes, a genetic diagnosis cannot be made^3^, suggesting as-yet-unidentified genetic cause(s).

β-cell glucose-stimulated insulin secretion (GSIS) is dependent on Ca^2+^ influx, through voltage-dependent calcium channels (VDCC)^5,6^. Reduced Ca^2+^ influx decreases GSIS; thus mutations that disrupt β-cell Ca^2+^ entry can cause MODY or the closely-related condition neonatal diabetes^7,8^. For example, gain-of-function mutations in K_ATP_ channel subunits hyperpolarize the β-cell membrane potential, reducing VDCC activity, Ca^2+^ influx and GSIS^7,8^. Other β-cell K^+^ channels, including two-pore domain K^+^ channels (K2P), also affect VDCC activity^9^. Expression of *KCNK16*, which encodes TWIK-related alkaline pH-activated K2P (TALK-1)^10^, is the most abundant and β-cell-selective of all human K^+^ channel transcripts^11,12^; and TALK-1 gain-of-function mutations would be predicted to cause diabetes similarly^9^.

Here we have used exome sequencing to identify the first family with MODY due to a mutation in *KCNK16*. The Leu114Pro substitution in TALK-1 affects the K^+^ selectivity filter, causing a profound increase in K^+^ current, altering β-cell Ca^2+^ flux, and decreasing GSIS in both human and mouse islet cells.

## METHODS

### Clinical Recruitment

A four-generation family with six affected family members with autosomal dominant diabetes was identified through a non-obese proband who presented aged 15 years (Fig.1A) with elevated fasting plasma glucose (7.8mmol/L) and an abnormal oral glucose tolerance test (glucose 19.6mmol/L two hours after 75g glucose). Antibody testing (islet cell, islet antigen 2, and glutamic acid decarboxylase-65) was negative. Over two decades, the proband required minimal insulin to maintain HbA1c of 5.7-6.5%; and she did not experience ketosis or other diabetes-related complications. Sanger sequencing for mutations in *GCK, HNF1A* and *HNF4A* (the commonest MODY genes) was negative. Other family members manifest diabetes similarly (detailed in Supplementary Appendix: Extended Clinical Data). The study protocol was approved by the relevant human research ethics committee (approval HREC/12/QPAH/109). All living family members gave written informed consent.

**Figure 1:**
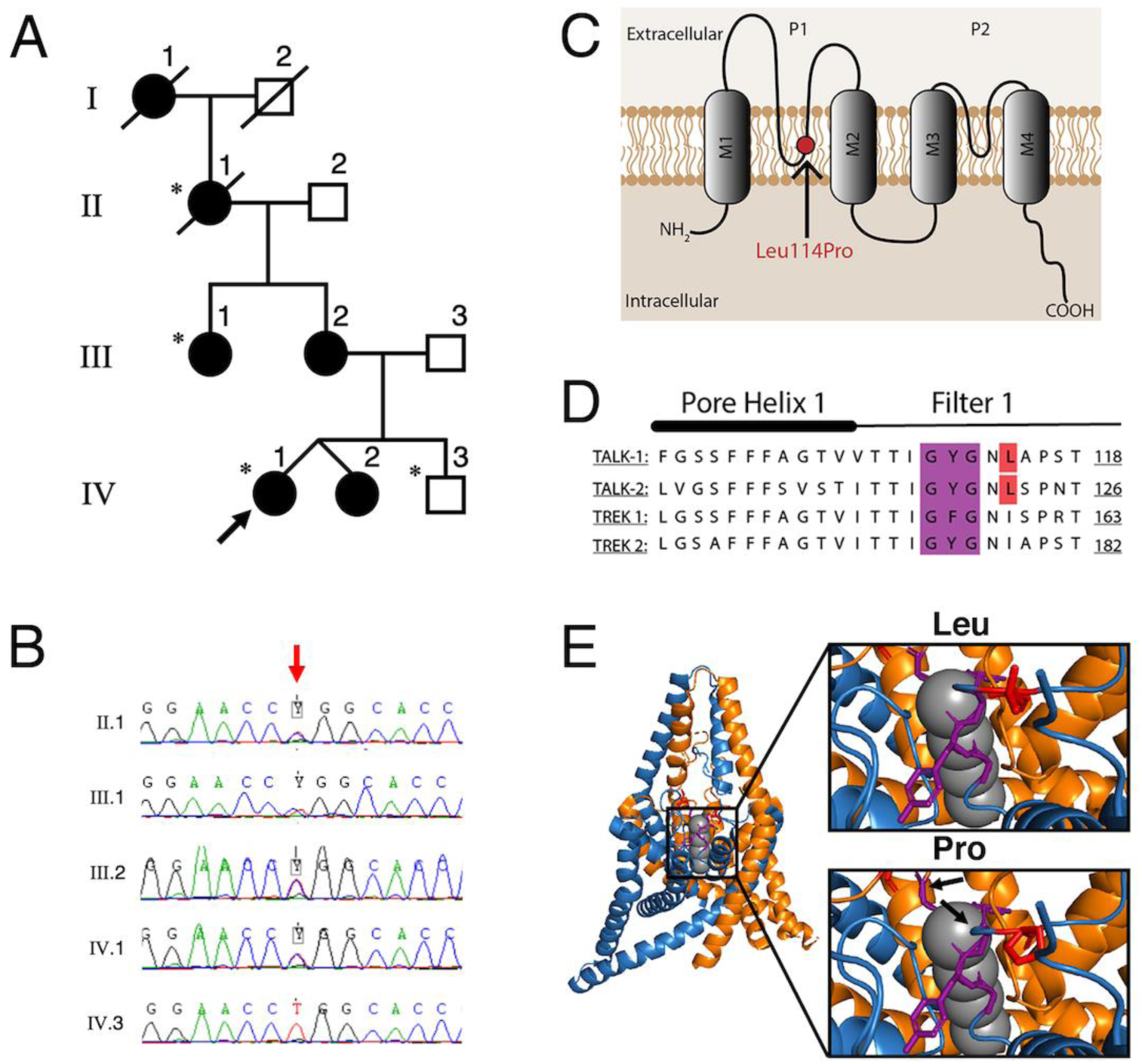
A novel *KCNK16* mutation co-segregates with MODY in a four-generation family and is predicted to affect the K^+^-selectivity channel of TALK-1. Family pedigree (Panel A: asterisks indicate individuals undergoing WES; filled-in shapes indicate individuals with diabetes; arrow indicates proband), with chromatogram of *KCNK16* variant (c.341T>C) (Panel B; red arrow indicates variant). Location of the predicted protein change (p.Leu114Pro), within the first pore domain and K^+^-selectivity channel of TALK-1 (Panel C), with alignment of Pore Helix 1 and Filter 1 amino acid sequences of *KCNK16* with other KCNK channels (Panel D; mutation position indicated in red; selectivity filter indicated in purple). Predicted conformational shifts (indicated by the arrows) in the K^+^ selectivity filter, modelled using TREK2 crystalline structure (Panel E).

### Exome sequencing

Exome sequencing, pipeline processing, quality control and variant curation was performed as previously described^13^ (detailed in Supplementary Appendix: Methods). Exome data from the proband was analysed for good-quality likely damaging rare variants in known MODY genes^1^, using a conservative minor allele frequency (MAF) threshold of <0.001, based on: (a) prevalence of paediatric diabetes of 0.2%^14^; and (b) prevalence of MODY mutations in 2% of a pediatric diabetes population^2^; further, most MODY mutations are private. Exome sequence data from the pedigree was analysed for novel and rare (MAF<0.001) good quality variants, of potentially damaging consequence, affecting highly conserved bases with appropriate segregation (i.e., heterozygous in affected individuals, absent in unaffected individuals).

### Plasmids and Transient Expression

Human TALK-1 wildtype (WT) and TALK-1 Leu114Pro constructs were created by site-directed mutagenesis and then cloned into a vector containing a P2A cleavage site followed by mCherry (Supplementary Appendix: Fig. S1). HEK293 cells, which have no endogenous TALK-1 expression, were transfected with 2µg DNA using Lipofectamine 3000 (Life Technologies). Transfection efficacy was assessed and quantified using mCherry Fluorescence. (Supplementary Appendix: Methods and Fig. S2).

### Lentivirus Production

HEK293 cells were transfected with lentiviral-producing plasmids; the plasmids used included the packaging plasmid (pCMV-dR7.74psPAX2), envelope plasmid (pMD2.G), and an expression plasmid (detailed in Supplementary Appendix: Methods and Fig. S1). Lentiviral-containing supernatants were collected three days after transfection and used for transduction of primary beta-β-cells; after transduction, equal TALK-1 expression was confirmed by equivalent mCherry expression (detailed in Supplementary Appendix: Methods and Fig. S3).

### Electrophysiological Recordings

TALK-1 channel currents were recorded in HEK293 cells using a whole-cell voltage-clamp technique with an Axopatch 200B amplifier and pCLAMP10 software (Molecular Devices), as previously described^9^; single-channel current recordings of TALK-1 were recorded with an cell-attached voltage-clamp technique also as previously described^15^ (detailed in Supplementary Appendix: Methods and Fig. S4).

### Islet and β-Cell Isolation

Islets were isolated from mouse pancreata as previously described^9^. Human islets from non-diabetic adult donors were provided by isolation centers of the Integrated Islet Distribution Program (donor information, Supplementary Appendix: Table S1). Some islets were dispersed into cell clusters and then cultured for 12–18 hours^9^. Cells were maintained in RPMI 1640 with 15% FBS, 100IU/mL penicillin, and 100mg/mL streptomycin in a humidified incubator at 37°C with an atmosphere of 95% air and 5% CO_2_.

### Calcium handling measurements

Islets were incubated for 25min in RPMI supplemented with Fura-2, AM (Molecular Probes), followed by incubation in Krebs-Ringer–HEPES buffer with 2mmol/L glucose for 20min^9^. For cytoplasmic Ca^2+^ [Ca^2+^_cyto_]: Ca^2+^ imaging was performed as previously described^9^, switching from 2mM glucose to 20mM glucose. For endoplasmic reticulum (ER) Ca^2+^ [Ca^2+^_ER_]: Islets were perfused in Krebs-Ringer–HEPES buffer without extracellular Ca^2+^ and 100 μM diazoxide and monitored for Ca^2+^_ER_ release mediated through blockade of the sarco(endo)plasmic reticulum Ca^2+^-ATPase (SERCA) with 50 μM cyclopiazonic acid (CPA, Alomone labs), as previously described^16^.

### Insulin Secretion Measurements

Islets were transduced with lentiviruses containing a RIP promoter^17^ upstream of either TALK-1 WT or TALK-1 Leu114Pro followed by a P2A cleavage site and NanoLuc-proinsulin (Supplementary Appendix: Fig. S1)^18,19^. Importantly, NanoLuc is co-secreted with insulin, enabling measurement of insulin secretion specifically from cells expressing either the TALK-1 WT or TALK-1 Leu114Pro construct^18^. Insulin secretion was measured as previously described^18^ (detailed in the Supplementary Appendix: Methods).

### Statistical Analyses

Functional data were analyzed using pCLAMP10 or Microsoft Excel and presented as mean ± SEM. Statistical significance was determined using Student’s *t*-test; a two-sided *P*-value ≤0.05 was considered statistically significant.

## RESULTS

### Exome sequencing in a family with MODY identifies a novel variant in KCNK16

Exome sequencing and analysis of known MODY genes in the proband identified a splice site mutation in *ABCC8* (NM_000352 c.1332+4 delC); however, this variant was not predicted to affect splicing^20^ and did not segregate appropriately in the pedigree.

Exome sequencing and analysis of the extended pedigree identified novel good-quality coding variants in two genes, *KCNK16* and *USP42*, with appropriate segregation (Fig. 1A and 1B; coverage statistics for exome sequencing, Supplementary Appendix: Table S2; exome data filtering, Supplementary Appendix: Table S3). *USP42* (*Ubiquitin-specific peptidase 42*) is involved in spermatogenesis^21^ and is not expressed in the pancreas; and was considered an unlikely MODY candidate. However, *KCNK16 (Potassium channel, subfamily K, member 16)* encodes for TALK-1, with its established role in GSIS^9^. Further, the *KCNK16*-containing locus is associated with T2DM^9,22,23^.

The *KCNK16* variant (NM_001135105: c.341T>C) has not previously been reported in ExAC (http://exac.broadinstitute.org), 1000 Genomes (http://www.1000genomes.org), or dbSNP137 (http://www.ncbi.nlm.nih.gov/projects/SNP/) databases. It affects a highly conserved base (GERP score 5.65) with the resultant amino acid change (p.Leu114Pro) predicted to involve the pore domain one of TALK-1, immediately downstream of the GYG K^+^ selectivity filter (Fig. 1C). The GYG motif, and leucine 114 specifically, shows strong sequence homology with other K2P channels (Fig. 1D). As the crystal structure of TALK-1 is unpublished, TREK-2 was used to model the p.Leu114Pro mutation which demonstrated a conformational shift in both the GYG motif and pore domain (Fig. 1E), strongly suggesting that TALK-1 Leu114Pro would significantly affect K^+^ permeability.

### TALK-1 Leu114Pro results in a gain-of-function

K^+^ currents recorded using HEK293 cells transfected with either TALK-1 WT or TALK-1 Leu114Pro demonstrated that TALK-1 Leu114Pro caused a massive (312-fold) increase in whole-cell K^+^ currents compared to TALK-1 WT (current at −40mV: TALK-1 Leu114Pro 774.16 ±218.75 pA vs. TALK-1 WT 2.48 ± 1.86 pA; Fig. 2A, 2B, Supplementary Appendix: Fig. S4). Individual TALK-1 channel activity showed a 3.6-fold increase in current amplitude and a 2.9-fold increase in open probability at 100mV for TALK-1 Leu114Pro compared to TALK-1 WT (Fig. 2C, 2D, 2E, 2F)^15^. Thus, TALK-1 Leu114Pro is a gain-of-function mutation, predicted to cause hyperpolarization of the β-cell membrane potential.

**Figure 2:**
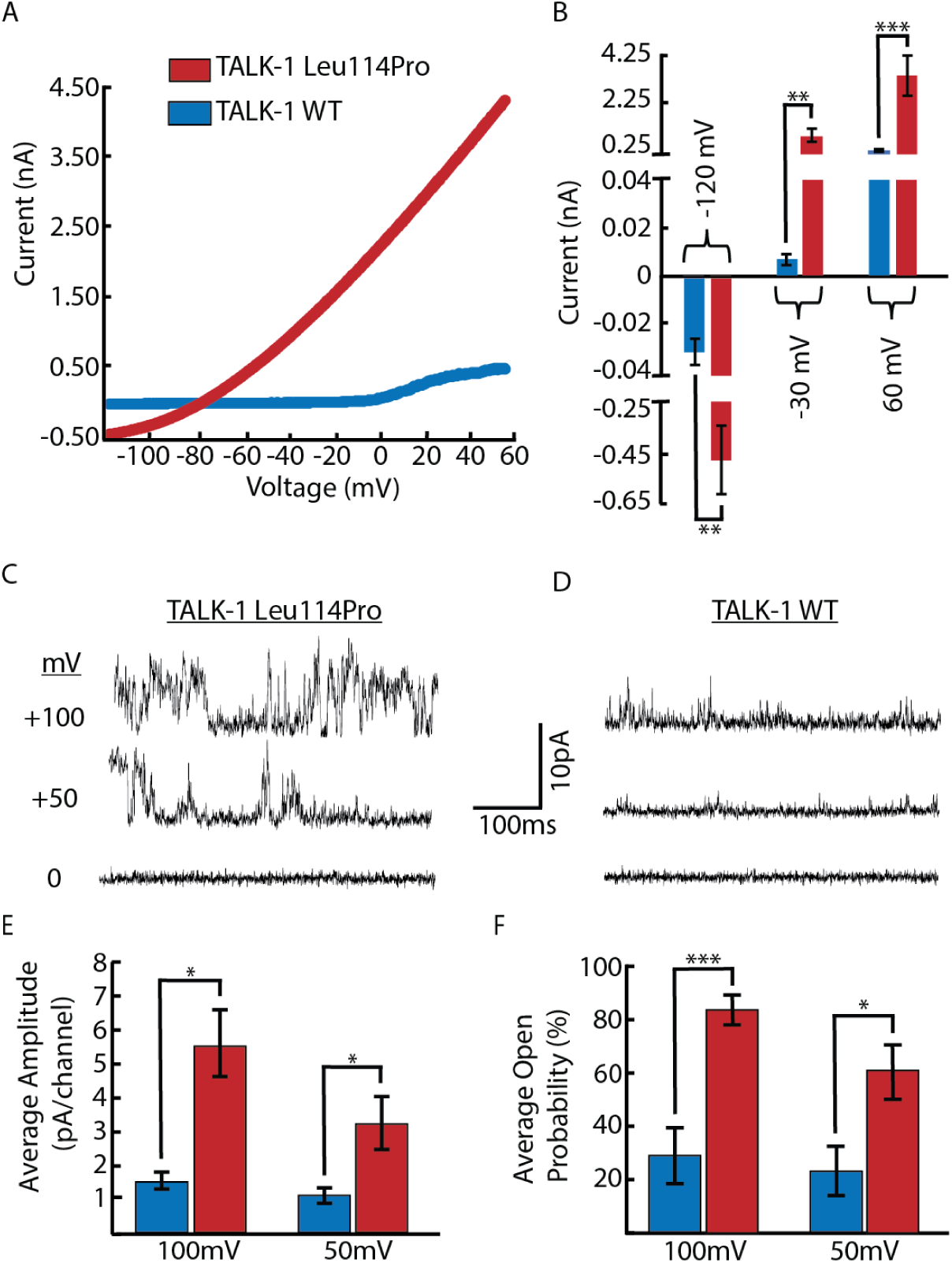
TALK-1 Leu114Pro causes a drastic gain-of-function in TALK-1 K^+^ current. K^+^ currents monitored from TALK-1 WT or TALK-1 Leu114Pro with whole-cell voltage clamp recordings, in response to a voltage ramp from −120mV to 60mV (Panel A); mean ± SEM; N= 11 control cells; N= 10 TALK-1 Leu114Pro cells (Panel B). Single-channel plasma membrane K^+^ currents monitored through TALK-1 Leu114Pro or TALK-1 WT with attached patch voltage clamp recordings, in response to the indicated voltage steps (Panel C and Panel D). Single channel recordings were analyzed for current amplitude (Panel E) and channel open probability (Panel F); mean ± SEM; N= 8 TALK-1 WT cells; N= 11 TALK-1 Leu114Pro cells \**P*<0.05, \*\**P*<0.01, \*\*\**P*<0.001.

### TALK-1 Leu114Pro reduces β−cell Ca^2+^ influx and ER Ca^2+^ stores

Ca^2+^ handling was monitored in mouse β-cells following transduction of either TALK-1 WT or TALK-1 Leu114Pro. Glucose-stimulated (20mM) β-cell Ca^2+^ influx was abolished by expression of TALK-1 Leu114Pro (Fig. 3A, 3C, and 3D). TALK-1 has been previously shown to modulate Ca^2+^_ER_ homeostasis by providing a countercurrent for Ca^2+^_ER_ release^16^; thus TALK-1 Leu114Pro control of Ca^2+^_ER_ storage was also examined. Inhibition of SERCAs with CPA resulted in significantly less Ca^2+^_ER_ release in β-cells expressing TALK-1 Leu114Pro compared to TALK-1 WT (62.6% decrease; Fig. 3E, 3F), suggesting reduced Ca^2+^_ER_ storage with TALK-1 Leu114Pro^16^. β-cells expressing TALK-1 Leu114Pro also showed elevated basal [Ca^2+^ _cyto_] compared to β-cells expressing TALK-1 WT (28.8% increase in AUC; Fig. 3A, 3B). Taken together, this suggests that under basal conditions, TALK-1 Leu114Pro enhances Ca^2+^_ER_ leak, thereby increasing basal [Ca^2+^_cyto_]. Furthermore, the usual transient drop in β-cell [Ca^2+^ _cyto_] following glucose stimulation of Ca^2+^_ER_ uptake (termed phase-0) was amplified with TALK-1 Leu114Pro compared to TALK-1 WT^24^. These changes would be predicted to diminish GSIS.

**Figure 3:**
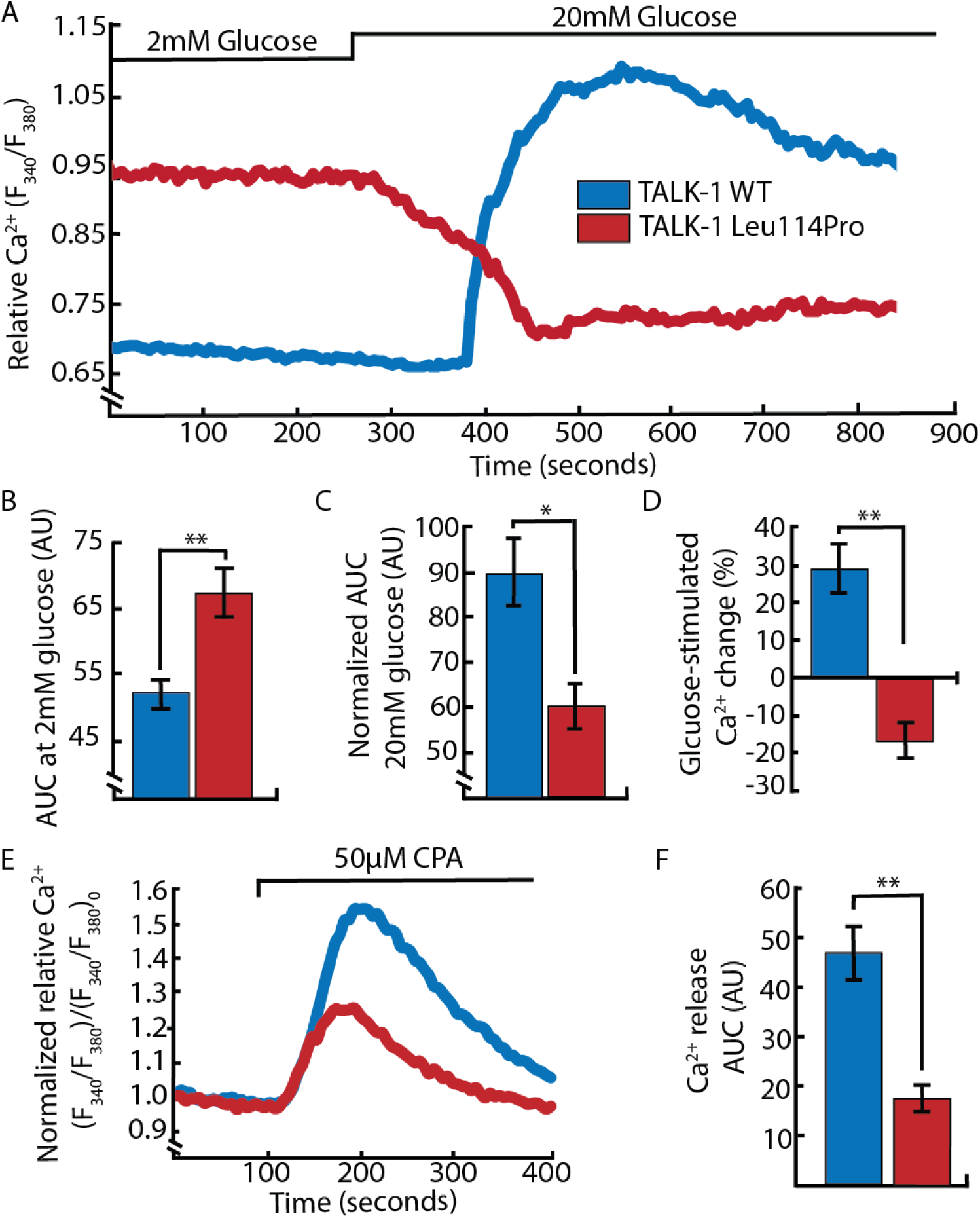
TALK-1 Leu114Pro modulates β-cell Ca^2+^ homeostasis. Representative β-cell Ca^2+^ measurements in response to 2mM and 20mM glucose (Panel A). Area under the curve (AUC) analysis of β-cell Ca^2+^ under low (2mM) glucose (Panel B) and high (20mM) glucose conditions (Panel C). AUC percent change from low glucose to high glucose (Panel D). Representative β-cell Ca^2+^ measurements in response to [Ca^2+^_ER_] depletion by CPA (Panel E) and the AUC analysis of the CPA response (Panel F). mean ± SEM; N= 3 animals for TALK-1 WT and TALK-1 Leu114Pro Ca^2+^ experiments except the for the 2mM condition that included N=9 animals. \**P*<0.05, \*\**P*<0.01, \*\*\**P*<0.001.

### TALK-1 Leu114Pro reduces Glucose-Stimulated Insulin Secretion

β-cells expressing TALK-1 Leu114Pro showed comparable basal (5mM glucose) insulin secretion but reduced GSIS (14mM glucose) compared to TALK-1 WT, in both mouse (52% decrease in GSIS) and human (38% decrease in GSIS) islets (Fig. 4).

**Figure 4:**
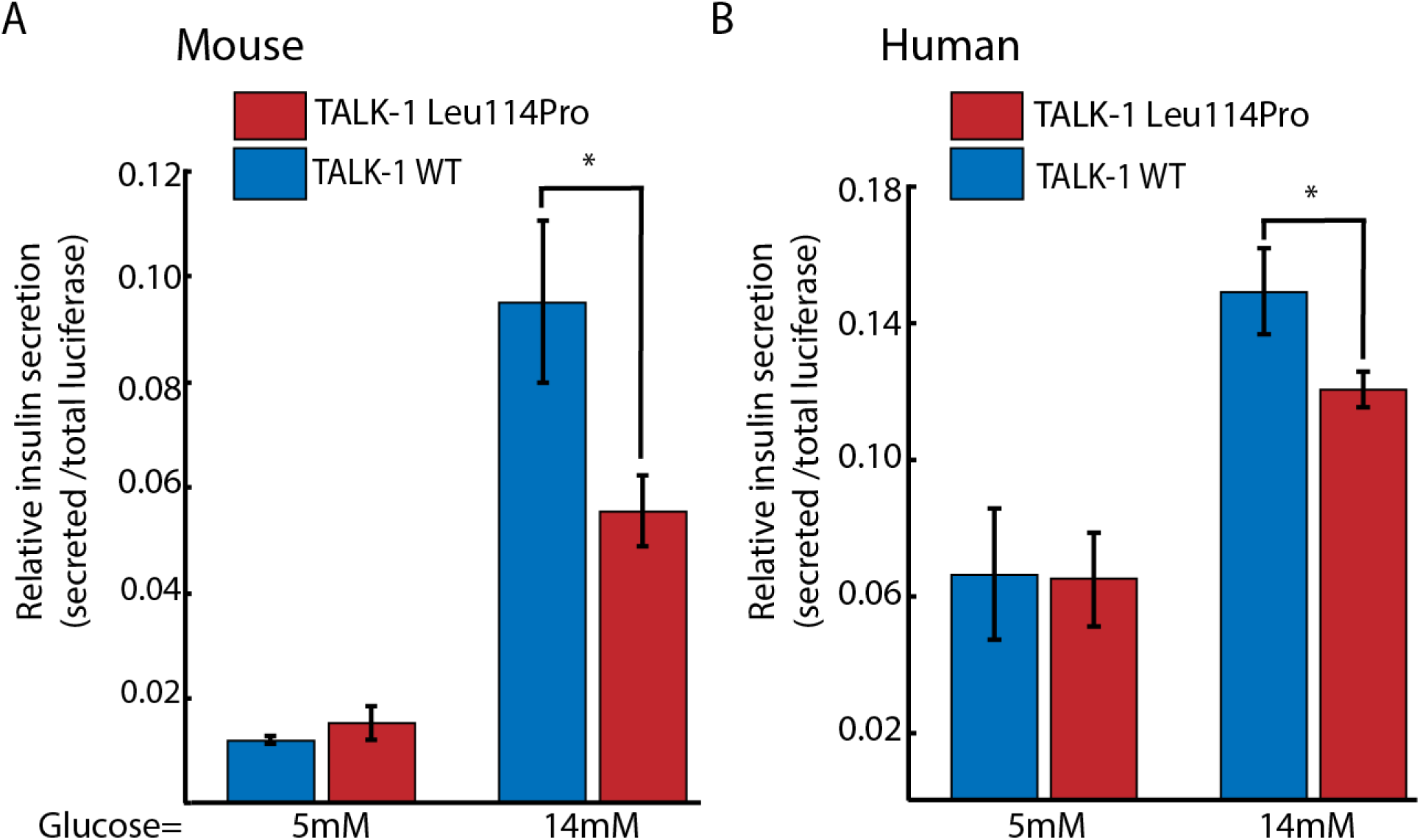
TALK-1 Leu114Pro reduces glucose-stimulated insulin secretion. Mouse (Panel A) and human (Panel B) islets transduced with viruses selectively expressing either TALK-1 WT or TALK-1 Leu114Pro and the NanoLuc-proinsulin luciferase insulin reporter. Islets were monitored for total secreted luciferase following exposure to 5mM or 14mM glucose; mean ± SEM; N=6 animals (14mM glucose, Panel A), N=3 animals (5mM glucose, Panel A); N=8 Human Donors (14mM glucose, Panel B), N=5 Human Donors (5mM glucose, Panel B). \**P*<0.05

## DISCUSSION

We have identified the first family with MODY due to a mutation in *KCNK16*. The novel TALK-1 gain-of-function p.Leu114Pro mutation increases β-cell K^+^ efflux resulting in membrane hyperpolarization, altering β-cell Ca^2+^ handling and decreasing GSIS; and highlight the critical role of TALK-1 in β−cell physiology. Unlike the only other MODY-associated K^+^ channelopathy (i.e., K_ATP_ channel dysfunction), TALK-1 is unresponsive to sulfonylureas^9^. Thus our data suggest not only a novel therapeutic target for *KCNK16*-associated MODY but for other forms of diabetes also.

TALK-1 belongs to the K2P channel family characterized by constitutive K^+^ flux, which serve critical roles setting the membrane potential of electrically excitable cells. The *KCNK16* transcript encoding TALK-1 is the most abundant K^+^ channel transcript in the human β-cell^11,12^; and *KCNK16* shows the most islet-selective expression of all ion channel transcripts^10,25^. Similar to other K^+^ channels, such as K_ATP_26, TALK-1-mediated hyperpolarization of mouse and human β-cell membrane potential limits VDCC activity, Ca^2+^ entry and GSIS^9^. However, the K_ATP_ K^+^ conductance is significantly greater than the small constitutive conductance of TALK-1^9,10,15,26^. Thus, TALK-1 mainly regulates the β-cell membrane potential following glucose stimulation, when K_ATP_ channels close: their activity limits islet Ca^2+^ oscillation frequency and hence GSIS^9^. A gain-of-function TALK-1 mutation would be predicted to affect glucose tolerance adversely resulting in hyperglycemia, as demonstrated here.

The *KCNK16*-containing locus is strongly associated with T2DM, in multiple genome-wide association studies including populations with differing ethnicities^22,23,27-29^, with strongest association (p<2×10^−8^) observed with the common nonsynonymous polymorphism rs1535500 (minor allele frequency 0.41, ExAC database, non-Finnish European descent subjects). The protein change (p.Ala277Glu) affects the C-terminal tail of TALK-1 and causes a modest (1.4-fold) increase in TALK-1 channel current, with both enhanced open probability and increased cell surface localization^9^. The risk haplotype is also associated with increased expression of the adjacent gene *KCNK17* which encodes another K2P channel, TALK-2^30^. TALK-2 is also expressed in islet cells with high specificity, though lower than TALK-1 (islet expression specificity index for *KCNK16*=0.98 and for *KCNK17*=0.76)^30^. These data suggest that the association of this locus with T2DM is driven by more than one mechanism, with combined effects from overactive TALK-1 and overexpression of TALK-2, each of which potentially contributing to hyperpolarization of the β-cell membrane potential, reducing glucose-stimulated Ca^2+^ influx and GSIS. That association is seen in T2DM with a common variant in the same gene in which we are now reporting a novel mutation causing MODY indicates that the encoded protein (i.e., TALK-1) is not functionally redundant; rather, that it is likely relevant to a high proportion of T2DM cases. These properties increase the potential of TALK-1 as a therapeutic target for T2DM.

Mutations in K2P channels causing dramatic changes in K^+^ channel currents typically affect the pore domains of these channels^31-33^. For example, loss-of-function mutations in the first or second pore domains of *KCNK3* (respectively, p.Gly97Arg and p.Gly203Asp) cause pulmonary hypertension^31^. Similarly, a loss-of-function mutation in the first pore domain of TASK-2 (p.Thr108Pro) causes Balkan Endemic Nephropathy^32^. A gain-of-function mutation (p.Gly88Arg) in the first pore domain of TALK-2, coded by *KCNK17*, causes a severe cardiac arrhythmia^33^; and is the only previously identified disease-associated mutation in TALK channels.

Gain-of-function mutations in K_ATP_ significantly increase β-cell K^+^ flux, resulting in neonatal diabetes and MODY^7,8^. In contrast, TALK-1 p.Leu114Pro results in a more modest diabetes phenotype despite the 300-fold increase in whole-cell TALK-1 activity. This may because TALK-1 activation shows voltage-dependence^9,10^. Unlike K_ATP_, which is active at all voltages, TALK-1 is an outward rectifying channel that shows increased activation during depolarization^9,10^. Therefore, a gain-of-function in TALK-1 would be most active post-β-cell depolarization – limiting, but not abrogating, insulin secretion. The p.Leu114Pro mutation does increase TALK-1 current near the resting membrane potential **(Fig. 2A, and Supplementary Appendix Fig. S4**); however, this current is still less than the total β-cell K_ATP_ conductance under euglycemic conditions. These data concord with the proband’s clinical phenotype, with dramatic glucose elevation after an oral glucose load but only a modest increase in fasting plasma glucose.

TALK-1 is expressed on both the β-cell plasma membrane and the ER membrane^16^. Ca^2+^_ER_ release is balanced by negative charge on the luminal ER membrane; this charge is dissipated by ER TALK-1 K^+^ influx leading to enhanced Ca^2+^_ER_ release^16^. Thus overactive TALK-1 channels (e.g., TALK-1 Ala277Glu) increase Ca^2+^_ER_ release, whereas TALK-1 ablation reduces Ca^2+^ release^16^. Importantly, enhanced β-cell Ca^2+^_ER_ release under hyperglycemic conditions results in ER-stress, contributing to β-cell dysfunction^34^. TALK-1 Leu114Pro may contribute to β-cell dysfunction via ER-stress, as observed in some MODY subtypes (e.g., *INS* mutations in MODY-10^35^); however, this remains speculative. Additionally, although highly β-cell specific, TALK-1 is also expressed in human pancreatic δ-cells where it negatively regulates somatostatin release ^36^. TALK-1 KO mice show increased somatostatin secretion under low and high glucose conditions, due to enhanced Ca^2+^_ER_ release^36^; thus, a gain-of-function mutation in TALK-1 may reduce δ-cell somatostatin secretion. The glycemic effects of this would be complex given the inhibitory effect of somatostatin on both insulin and glucagon secretion^36^; and require future investigation.

Some MODY subtypes (e.g., *ABCC8-, KCNJ11-, HNF1α*- and *HNF4α* MODY) are manageable through K_ATP_ inhibition^7,37^,38 – i.e., sulfonylurea use. Although β-cell membrane potential depolarization with sulfonylureas may allow greater VDCC activity, potentially increasing insulin secretion in affected individuals in this family, TALK-1 itself is not sensitive to sulfonylureas^9^. Further, and as detailed above, TALK-1 primarily modulates β-cell membrane potential during active insulin secretion when K_ATP_ is closed (i.e., during hyperglycemic conditions)^9^. Thus, K_ATP_ inhibition may not completely normalize β-cell membrane potential or insulin secretion in individuals with TALK-1 gain-of-function MODY. This raises the possibility of TALK-1 inhibition as a druggable target. Genetic evidence whether from rare (e.g., MODY) or common (e.g., T2DM^22,23,27-29^) human disease is a strong predictor of future successful drug development^39^. Thus our data have important therapeutic implications not only for TALK-1 MODY but also for the far more common form of diabetes T2DM.

In conclusion, we have identified a novel mutation in *KCNK16* causing a gain-of-function in TALK-1, reducing glucose-stimulated Ca^2+^ influx, Ca^2+^_ER_ storage, and GSIS; and resulting in MODY. TALK-1 is the first ion channel linked to MODY after K_ATP_; and is expressed more selectively in islet cells compared to K_ATP_. The *KCNK16* locus is associated with T2DM risk in the general population. Our data suggest TALK-1 as an efficacious and islet-selective therapeutic target for both *KCNK16*-associated MODY and T2DM.

## Acknowledgments

The authors gratefully acknowledge the laboratory support of Mr Lawrie Wheeler, Ms Lisa Anderson, Ms Sharon Song, and Ms Jessica Harris; and the editorial support of Dr David Pennisi. Sarah Graff was supported by a Ruth L. Kirschstein National Research Service Award (NRSA) Individual Predoctoral Fellowship (F31 DK118855). Dr Stephanie Wilson was supported by a University of Queensland Research Scholarship. Dr Aideen McInerney-Leo was funded by a National Health and Medical Research Council (NHMRC) Early Career Fellowship (ID 1158111). Prof Matthew Brown was supported by an NHMRC Senior Principal Research Fellowship (ID 1024879). The Translational Research Institute was supported by a grant from the Australian Government. This project was supported by a grant from the Royal Brisbane and Women’s Hospital Project Grant (2011) and an Australian Paediatric Endocrine Group research grant (2015). This work was also supported by grants (R01 DK-081666, R01 DK115620) from the National Institutes of Health, the American Diabetes Association (Grant 1-17-IBS-024), and a Vanderbilt University Diabetes Research Training Center Pilot and Feasibility Grant (P60-DK-20593).

Mrs Sarah Graff and Dr Stephanie Johnson contributed equally to this article. Dr Jacobson and Prof Duncan also contributed equally to this article.

