## Supplemental Appendix for "A novel mutation in *KCNK16* causing a gain-of-function in the TALK-1 potassium channel: a new cause of maturity onset diabetes of the young"

**Supplementary Appendix**


| **Table of Contents** | **Page Number** |
|---|---|
| Supplementary Methods | 2 |
| Supplementary Results | 6-14 |
| • Clinical Data | 6 |
| • Table S1 | 7 |
| • Table S2 | 8 |
| • Table S3 | 9 |
| • Figure S1 | 10 |
| • Figure S2 | 11 |
| • Figure S3 | 12 |
| • Figure S4 | 13 |
| Supplementary References | 14 |

### Supplementary Methods

#### *Exome sequencing and analysis*

DNA was extracted from blood or saliva and sequencing libraries constructed according to each manufacturer's instructions. Exome capture was performed using either Nimblegen SeqCap EZ Human v3.0 Exome Enrichment Kit for the proband (Roche, Basel, Switzerland) or Nextera™ Rapid Capture Exome kit for subsequent family members (Illumina, San Diego, CA, USA). Pre- and post-capture quality and yield were assessed by Agilent High Sensitivity DNA assay (Agilent, Santa Clare, CA) using Bioanalyzer 2100 (Agilent, Santa Clare, CA) and by qPCR using the KAPA Library Quantification Kit (Kapa Biosystems, Wilmington, MA).

Multiplexed massively parallel sequencing (six samples per flowcell lane) was performed on an Illumina HiSeq 2000 for the proband or HiSeq 2500 for additional family members (Illumina, San Diego, CA, USA), generating 100-base pair paired-end reads.

Data were demultiplexed using Illumina Data Analysis Pipeline software (Bcl2fastq v1.8.4) and aligned to the current reference human genome (hg19, released February 2009) using the Novoalign alignment tool (V3.00.02). Sequence alignment files were converted using Picard tools (v1.124). Variants were called using the Genome Analysis Toolkit (GATK v2.7-2) and annotated using ANNOVAR. Further analysis of sequence data was performed using custom scripts employing R and Bioconductor.

After quality control and exclusion of artefact, good quality variants with minor allele frequency (MAF) <0.05 (assessed against internal and external databases, including ExAC[1], 1000 Genomes[2], and dbSNP137[3]) were retained for analysis.

#### *Lentiviral Plasmids*

Lentiviral plasmids expressing TALK-1 WT and TALK-1 Leu114Pro were designed for HEK cell and β-cell studies. A cartoon representation of the different lentiviral constructs that were utilized in these experiments is presented in **Supplementary Figure S1**. For whole-cell current recordings, a lentiviral plasmid was created that contains the CMV enhancer and CMV promoter placed immediately upstream of either TALK-1 WT or TALK-1 Leu114Pro followed by a P2A cleavage site and mCherry. For the β-cell $Ca^{2+}$ experiments, a lentiviral plasmid was created that contains the β-cell specific RIP1-mini-CMV promoter[4] placed immediately upstream of either TALK-1 WT or TALK-1 Leu114Pro. To utilize the Nano-

Luc Pro-insulin Luciferase[4] as a read-out of insulin secretion we developed lentiviral plasmids that contained the same RIP1-mini-CMV Promoter mentioned above expressing either TALK-1 WT or TALK-1 Leu114Pro followed by a P2A cleavage site and the Nano-Luc Pro-insulin Luciferase. A CMV promoter expressing GFP was also inserted into the plasmid to enable visualisation of plasmid expression. The sequences for TALK-1 are those corresponding to transcript variant 3 (NM_001135105). Transcript Variant 3 was used for these experiments because it is the most highly expressed ion channel producing variant of *KCNK16* in human β-cells.

*Lentivirus Production*

HEK293FT cells were cultured in DMEM GlutaMax with 10% FBS, 100IU/mL penicillin, and 100mg/mL streptomycin to 50-70% confluency in 100mm dishes. The media was then switched to DMEM GlutaMax with 10% Heat-Inactivated FBS for transfection. Chloroquine was added to the cells and allowed to sit for 10 minutes. The following DNA mixture (46.7ul of 2.5M $CaCl_2$, 18.65ug lentiviral plasmid, 13.9ug packaging plasmid (pCMV-dR7.74psPAX2), and 5.56ug envelope plasmid (pMD2.G) brought up to 470ul with water) was added to 470ul of 2X HBS and then added dropwise onto the cells. Cells were incubated with DNA mixture for 5-7 hours and then the transfection media was aspirated off and replaced with fresh DMEM GlutaMax with 10% FBS, 100IU/mL penicillin, and 100mg/mL streptomycin. Media containing lentivirus was harvested 72 hours later and frozen at −80°C before use in mouse and human islet cells.

*Lentiviral Transduction*

For viral transduction, islets were isolated from mouse pancreata and dispersed into clusters through titration in 0.005% trypsin. After overnight recovery, the islet clusters were transduced using polybrene with 3$^{rd}$-generation lentiviruses containing the above-mentioned lentiviral plasmids. The virus was left on the cells for 5-7 hours and then replaced with fresh media. The virus was then allowed to express for 2-3 days. The mCherry fluorescence allowed us to identify β-cells that were successfully transduced and only mCherry positive cells were selected for analysis. On average, the mCherry fluorescence, and therefore expression level, was similar between TALK-1 WT and TALK-1 Leu114Pro expressing cells **(Supplementary Figure S3)**.

*TALK-1 Whole-Cell Currents*

Voltage-clamp mode on an Axopatch 200B amplifier (Molecular Devices) was used to measure whole-cell TALK-1 currents. A Digidata 1440 was used to digitize currents that were low-pass–filtered at 1 kHz. To record whole cell TALK-1 currents in HEK cells, a lentiviral plasmid was produced containing a CMV promoter expressing TALK-1 WT followed by a P2A cleavage site and mCherry. HEK cells were transfected with 2ug of the lentiviral plasmid DNA using Lipofectamine 3000. The mCherry allowed us to identify cells that were successfully transfected and currents were recorded from cells with similar levels of mCherry fluorescence. On average, the mCherry fluorescence, and therefore expression level, was similar between TALK-1 WT and TALK-1 Leu114Pro expressing cells **(Supplementary Figure S2)**. The whole cell currents were analysed using ClampFit and Excel.

*TALK-1 Single Channel Currents*

Single Channel currents were measured in HEK cells using attached-patch voltage-clamp technique. Electrodes were pulled to a resistance of 8 to 10 megaohms and then coated with Sigmacote. Extracellular solution contained 135mM NaCl, 5mM KCl, 1mM $MgCl_2$, 1mM $CaCl_2$, and 10mM Hepes (pH 7.3 with NaOH). Intracellular pipette solution contained 150mM KCl, 1mM $MgCl_2$, 5mM EGTA, and 10mM HEPES (pH 7.3 with KOH). Single channel current openings were analysed for open probability and current amplitude during a 5-second period of stimulation with 100mV, 50mV, and 0mV. A threshold of 0.75pA was set to determine channel openings. Our reported amplitude and open probability for TALK-1 WT is similar to that previously published for TALK-1[5,6].

*Calcium Imaging*

Islets were incubated at 37°C for 25min in RPMI supplemented with 2µmol/L Fura-2, AM (Molecular Probes), followed by incubation in Krebs-Ringer–HEPES buffer with 2mmol/L glucose for 20min. Cells were then imaged every 5 second using a Nikon Eclipse Ti2 microscope equipped with a Photometrics Prime 95B 25mm sCMOS Camera. The ratios of Fura-2AM fluorescence excited at 340 and 380 nm (F340/F380) were measured. The cells were perifused at a flow rate of 2 mL/min at 37°C with solutions containing glucose concentrations specified in the figures.

*Nano-Luc Pro-insulin Luciferase Assay*

Islet clusters were cultured in a 96-well plate at 20 islet equivalents per well. Islet clusters were transduced as described above with lentiviral plasmids that contained the RIP1-mini-CMV Promoter expressing either TALK-1 WT or TALK-1 Leu114Pro followed by a P2A cleavage site and the Nano-Luc Pro-insulin Luciferase. The islet clusters were starved for one hour in 100ul media (DMEM) containing 5mM glucose before running the assay. The assay was then run using the protocol for Promega Nano-Glo® Luciferase Assay System. In short, the starvation media is replaced with fresh media containing 5mM glucose and the islet clusters are allowed to secrete for one hour. The media is then collected and replaced with media containing 14mM glucose and the islet clusters are again allowed to secrete for one hour. After the secretion the cells are lysed and collected. The total lysate, the 5mM glucose secretion media, and 14mM glucose secretion media are mixed with the Promega Nano-Glo® substrate and imaged for luminescence using a BioTek Synergy™ H4 Hybrid Microplate Reader. Luminescence from the low and high glucose secretion media were divided by total luminescence from the lysate.

*Human Islet Cells*

Human islets were obtained for the Nano-Luc Pro-insulin Luciferase studies from the Integrated Islet Distribution Program (IIDP). Human donors were both male and female as well as from multiple ethnic backgrounds. Information on each human donor is presented in Supplementary Table S1.

**Supplementary Results**

*Extended Clinical Data*

The non-obese proband (individual IV.1) was diagnosed with antibody-negative diabetes age 15 years. Now age 32 years she only requires a maximum of 1-2 units of basal insulin to maintain an HbA1c 5.7-6.5%. She has not had ketosis or other diabetes-related complications.

Sanger sequencing screening for mutations *GCK, HNF1A* and *HNF4A* was negative.

The proband's identical twin sister (individual IV.2) was diagnosed with diabetes contemporaneously with the proband. This individual is managed with multiple daily injections of insulin. She has never experienced ketosis or diabetes-related complications. Other clinical information for this individual is unavailable.

The proband's mother (individual III.2) and aunt (individual III.2), both of normal weight, presented aged 15 and 13 years respectively with antibody-negative diabetes. Neither has experienced ketosis or diabetes-related complications after many decades. The proband's mother requires 8-11 units isophane insulin daily, maintaining an HbA1c of 6-7%. Her aunt requires 9-10 units of mixed insulin (Mixtard® 30/70 mane), maintaining an HbA1c <6%.

The proband's maternal grandmother (individual II.1) was diagnosed with diabetes during pregnancy, aged 30 years. She required <10 units/day of basal insulin, maintaining an HbA1c 6-6.5%. She recently died aged 82 years, without experiencing ketosis or diabetes-related complications.

The proband's maternal great-grandmother (individual I.1) also had diabetes. Further details of this individual are not available.

The proband's brother (individual IV.3), currently aged 26 years, does not have diabetes.

**Supplementary Tables and Figures**

**Table S1. Human islet donor information**. Human islets received for human insulin secretion studies from the Integrated Islet Distribution Program.

| Islet Isolation Center | Age | Gender | Ethnicity | Weight (kg) | Height (cm) | BMI | Cause of Death | Purity (%) | Average blood glucose in mmol/L |
|---|---|---|---|---|---|---|---|---|---|
| Southern California | 53 | Male | White | 83.9 | 182.9 | 25.1 | Stroke | 96 | 6.2 |
| Southern California | 51 | Male | White | 76 | 177.8 | 24.0 | Head Trauma | 96 | 7.8 |
| Miami | 57 | Female | Black | 79.5 | 165.0 | 29.2 | Stroke | 95 | 9.6 |
| Scharp/Lacy | 50 | Male | Hispanic | 83.9 | 160.0 | 32.8 | Anoxia | 90 | 7.5 |
| Scharp/Lacy | 57 | Male | Hispanic | 72.6 | 167.6 | 25.8 | Stroke | 95 | 7.8 |
| Wisconsin | 53 | Female | White | 38.1 | 145.0 | 40.0 | Stroke | 92 | 7.0 |
| Scharp/Lacy | 45 | Male | Black | 106.1 | 180.3 | 32.6 | Anoxia | 95 | 7.3 |
| Southern California | 33 | Male | White | 106.4 | 179.0 | 33.2 | Anoxia | 80 | 7.8 |

*1 mg/dL = 0.0555 mmol/L

**Table S2 Coverage statistics for exome sequencing**

| | MODY_II.1 | MODY_III.1 | MODY_IV.1 | MODY_IV.3 |
|---|---|---|---|---|
| **Total bases** | 5835073129 | 5386986408 | 5187196181 | 1824158333 |
| **% bases on target exons** | 38.4 | 35.9 | 47.9 | 35.8 |
| **Target exon median base coverage** | 52.9 | 44.2 | 32.0 | 12.8 |
| **Target exon mean base coverage** | 64.4 | 53.5 | 37.6 | 14.2 |
| **% target exon bases with coverage >1x** | 98.2 | 98.6 | 91.0 | 95.5 |
| **% target exon bases with coverage >5x** | 93.6 | 94.5 | 94.4 | 90.4 |
| **% target exon bases with coverage >10x** | 88.0 | 88.7 | 79.2 | 80.1 |
| **% target exon bases with coverage >30x** | 65.9 | 63.0 | 54.6 | 59.9 |

**Table S3: Bioinformatic filtering of identified variants**

| Filtering steps | MODY-II.1* | MODY-III.1* | MODY-IV.1* | MODY-IV.3 |
|---|---|---|---|---|
| Variants identified with MAF≤0.05 | 7114 | 7389 | 8368 | 7320 |
| Remaining variants of potentially damaging consequence** of good quality after removal of platform-related artefact | 659 | 639 | 434 | (NA) |
| Heterozygous variants remaining with MAF <0.001 | 343 | 333 | 213 | (NA) |
| Remaining variants segregating appropriately with observed autosomal dominant inheritance | 13 | | | |
| Remaining variants with high conservation (GERP >2.5) | 6 | | | |
| Remaining variants predicted to be damaging#; MAF reported from ExAC | *KCNK16 (*NM_001135105) c.341T>C; p.Leu114Pro; MAF = 0<br>*USP42* (NM_032172) c.C3569G: p.Pro1190Arg; MAF=0<br>*KIAA1407* (NM_020817) c.2687G>A; p.Arg89Gln; MAF 0.00007 (rs144192553) | | | |
| Remaining variants in candidate genes known to be involved in insulin secretion | *KCNK16 (*NM_001135105) c.341T>C; p.Leu114Pro | | | |

*affected individuals

**potentially damaging consquences defined as nonsynonymous single nucleotide variants (SNVs), splice site SNVs, frameshift substitution, stopgain or stoploss SNVs.

#Using protein prediction algorithms SIFT[7], Mutation Taster Human Splicing Finder v3.0[7] and PolyPhen2[8]

MAF: minor allele frequency. ExAC: Exome Aggregation Consortium. QC: quality control. GERP: genomic evolutionary rate profiling score.

**Figure S1. Lentiviral plasmids designed for TALK-1 WT and TALK-1 Leu114Pro expression in HEK cells and β-cells.**

Lentiviral Plasmids were designed with either a CMV promoter or a Rat-Insulin Promoter expressing TALK-1 followed by a P2A cleavage site and either mCherry (for electrophysiology a calcium experiments) or Pro-insulin Luciferase (For insulin secretion experiments). The TALK-1 cloned into these constructs were either Wild-Type (WT) **(Panel A)** or TALK-1 Leu114Pro **(Panel B).** Plasmids were sequenced and chromatograms show the correct TALK-1 WT **(Panel C)** and TALK-1 Leu114Pro **(Panel D)** sequences.

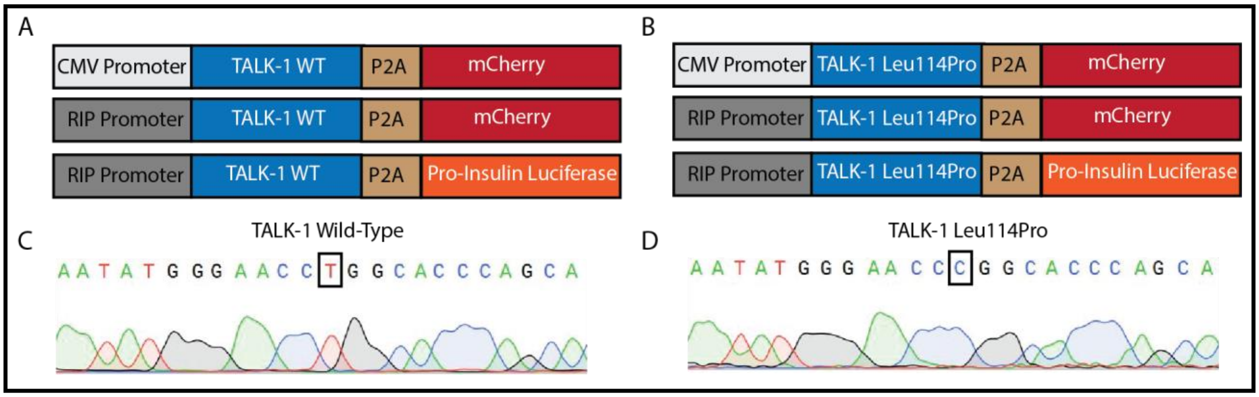

**Figure S2. Expression of TALK-1 WT and TALK-1 Leu114Pro in HEK cells for current recordings.**

HEK cells were transfected with constructs containing a CMV Promoter expressing TALK-1 followed by a P2A cleavage site and mCherry. TALK-1 expressing cells are represented by mCherry fluorescence (red) and all nuclei were stained with Hoescht (blue); cells expressing TALK-1 WT **(Panel A)** or TALK-1 Leu114Pro **(Panel B)** (images shown were taken at 20X magnification). Equivalent mCherry fluorescence in TALK-1 WT- and TALK-1 Leu114Pro-expressing cells indicates similar TALK-1 expression levels **(Panel C)** (N=30 for cells expressing TALK-1 WT; N=26 for cells expressing TALK-1 Leu114Pro).

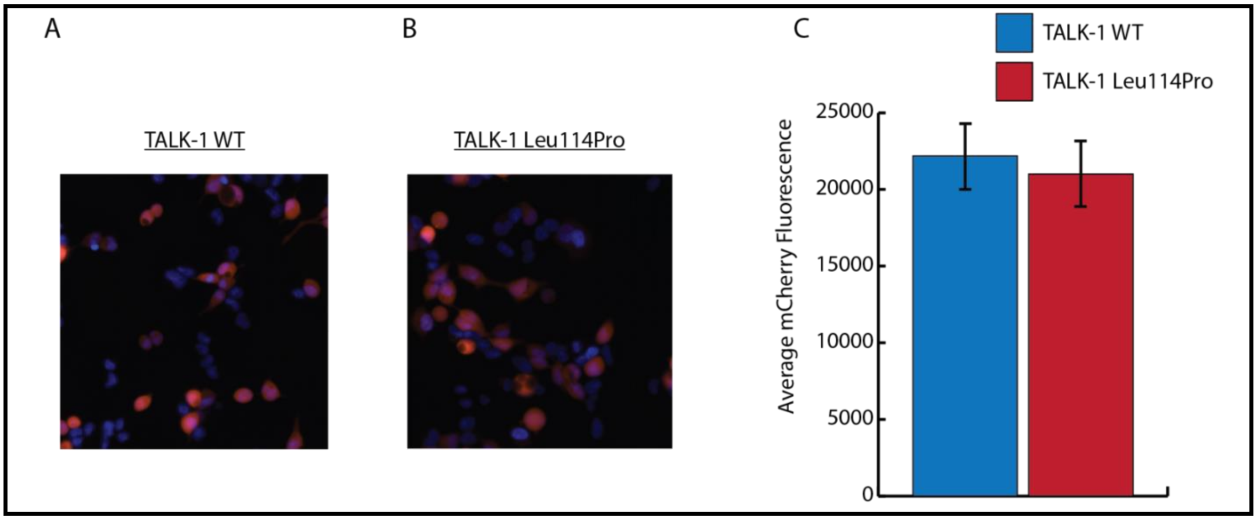

**Figure S3. Expression of TALK-1 WT and TALK-1 Leu114Pro in mouse β-cells.**

Dispersed islet cells were transduced with the viruses containing a Rat-Insulin Promoter expressing TALK-1 followed by a P2A cleavage site and mCherry. TALK-1 expressing β-cells (shown in red) imaged with a 10X objective compared to total islet cells (marked in blue) for TALK-1 WT **(Panel A)** and TALK-1 Leu114Pro **(Panel B)**. Average mCherry fluorescence shows similar expression levels between TALK-1 WT and TALK-1 Leu114Pro **(Panel C)** (N=48 for cells expressing TALK-1 WT; N=42 for cells expressing TALK-1 Leu114Pro).

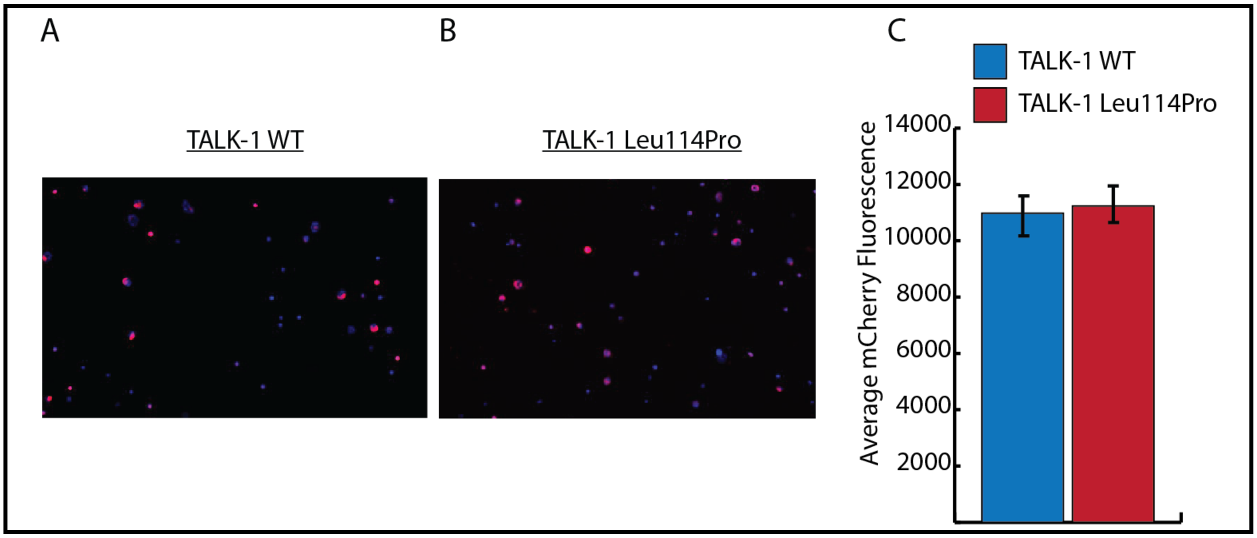

**Figure S4. Average whole cell TALK-1 current recordings in HEK cells expressing TALK-1 Leu114Pro vs. TALK-1 WT.**

Average whole cell currents from TALK-1 Leu114Pro-expressing cells (in red) is dramatically greater for TALK-1 WT (in blue) **(Panel A)**. A zoomed-in graph of the average TALK-1 WT whole-cell current shows K2P current similar to previously published TALK-1 WT currents[9] **(Panel B)**.

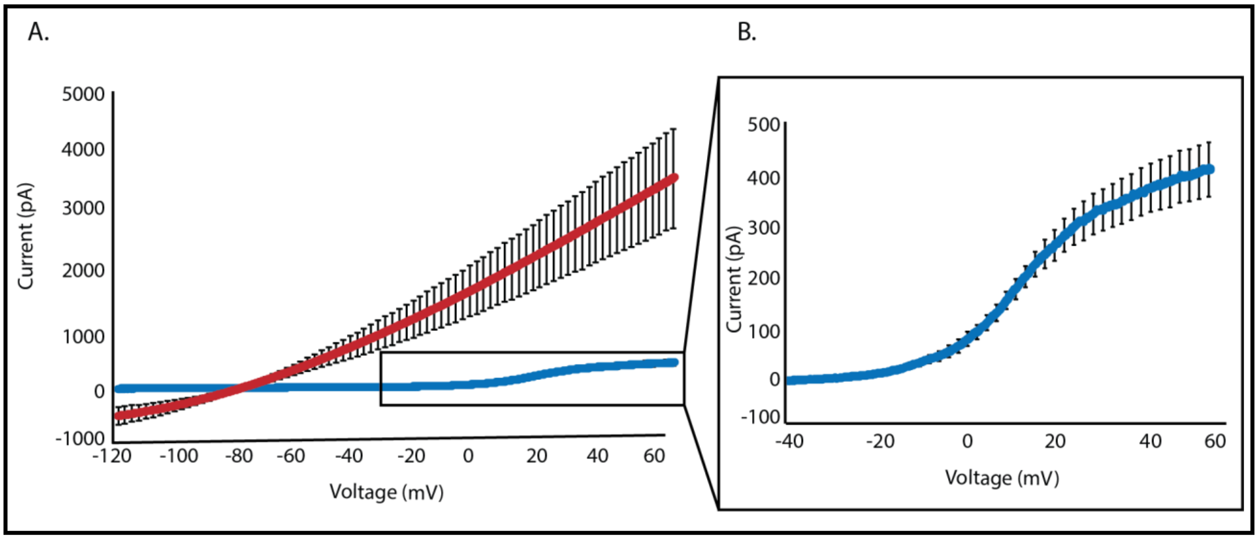
